# Structural Components for Calcitonin Gene-Related Peptide Signaling to Oligodendrocyte Precursor Cells

**DOI:** 10.64898/2026.03.23.713636

**Authors:** Rebecca Aitken, Yadong Ji, Thomas Blanpied, Asaf Keller, Rebecca Lorsung

## Abstract

Oligodendrocyte precursor cells (OPCs) are unique glial cells that communicate bidirectionally with neurons. Neuronal inputs drive various OPC behaviors, including proliferation and differentiation, immunomodulation, blood brain barrier regulation, synapse engulfment and axonal remodeling. OPCs are implicated in numerous stress and pain conditions, where their involvement is likely driven by neuronal activity (ie. neurotransmitter and neuropeptide signaling). One neuropeptide causally involved in chronic pain and stress conditions is calcitonin gene-related peptide (CGRP). Here, we tested the hypothesis that OPCs receive direct inputs from CGRP-containing neurons in the adult brain. Using RNAscope, immunofluorescence and analysis of single-cell datasets, we find that OPCs express receptors for CGRP and we identify close spatial contacts between CGRP and OPCs, with nearly half of CGRP puncta occurring within 1 µm of an OPC. Some of these contacts appear to be synaptic, with CGRP-OPC contacts colocalizing with the presynaptic protein Bassoon and the postsynaptic protein PSD-95. This work suggests the presence of both diffuse and more direct forms of CGRP signaling to OPCs, raising the importance of future experiments to identify both the mode of CGRP release onto OPCs and the functional effects of these different contact types.

## Introduction

Oligodendrocyte precursor cells (OPCs) are unique glial cells that contribute to neural function and dysfunction through a variety of actions. In both the developing and adult brain, OPCs differentiate into myelin-forming oligodendrocytes (Nishiyama et al., 2021; Young et al., 2013). However, a population of “adult” OPCs remains that is involved in additional functions, including synapse engulfment, axonal remodeling, blood-brain barrier regulation, and immunomodulation (Buchanan et al., 2023; Fang & Bai, 2023).

OPCs receive diverse neuronal inputs that affect their activity. Notably, OPCs are the only glial cell type that receive direct synaptic input from neurons (Bergles et al., 2000, 2010; Lin et al., 2005). In addition to direct glutamatergic and GABAergic synaptic input, suggested to drive OPC differentiation and myelination (J. Li et al., 2024), OPCs also respond to adrenergic and peptidergic signals. OPCs express a variety of G protein-coupled receptors, including those for dynorphin, norepinephrine (NE) and pituitary adenylate-cyclase activating protein (PACAP) (Fiore et al., 2023; Lee et al., 2001; Lu et al., 2023; Mei et al., 2016; Osso et al., 2021). Dynorphin release from neurons promotes OPC differentiation and myelination, while NE and PACAP increase OPC proliferation (Lee et al., 2001; Lu et al., 2023; Mei et al., 2016; Osso et al., 2021).

Dysfunctions of OPCs are implicated in several conditions, including mood and pain disorders. For example, in patients with depressive disorders, OPCs are one of the most differentially regulated cell types (Nagy et al., 2020). Activation of OPCs results in stress-related behaviors (X. Zhang et al., 2021), while chronic stress initially reduces, then increases OPC density in the hippocampus and prefrontal cortex (Birey et al., 2019). OPC proliferation increases in several animal models of chronic pain (Echeverry et al., 2008; Kim & Angulo, 2025). However, it is unknown which neuronal inputs to OPCs may influence their involvement in pain and stress disorders.

Chronic pain and anxiety disorders are comorbid, and share underlying biological mechanisms (Keller, 2020, 2023). A key component of these mechanisms is calcitonin gene-related peptide (CGRP), a neuropeptide with both peripheral and central nervous system roles. CGRP modulates several cardiovascular, neural, immune, and metabolic processes (Kee et al., 2018; Russo & Hay, 2023), and has established roles in the pathogenesis of chronic pain conditions, including migraine (Goadsby et al., 2017; Schou et al., 2017).

Previous studies—including from our laboratory—investigating the role of CGRP in chronic pain and stress-related disorders have assumed that its effects on the central nervous system are mediated by neuronal CGRP receptors (Lorsung et al., 2025; Smith et al., 2023). However, it remains unclear whether CGRP may directly influence glia, such as OPCs. Here, we tested the hypothesis that OPCs receive direct inputs from CGRP-containing neurons in the adult brain.

We focus particularly on the parabrachial nucleus (PBN) and its projection targets, because it is a main CGRP network in the brain (Huang et al., 2021; Palmiter, 2018) and because PBN dysfunctions are causally related to chronic pain and affective disorders (Palmiter, 2024; Raver et al., 2020).

## Materials and Methods

### Animals

All procedures adhered to the Guide for the Care and Use of Laboratory Animals and were approved by the Institutional Animal Care and Use Committee at the University of Maryland School of Medicine. We used CGRP^Cre^ heterozygous adult male and female mice that were bred in-house from male B6.Cg-Calca^tm1.1(cre/EGFP)Rpa^/J (strain #033168) × female C57BL/6J mice (strain #000664). Breeding pairs were obtained from The Jackson Laboratory. Offspring were weaned at postnatal day (P)21 and housed two to five per cage in single-sex groups. Food and water were available *ad libitum*, and lights were maintained on a 12 h light/dark cycle.

Mice between 6-8 weeks old were anesthetized with an intraperitoneal injection of ketamine (1.5µg/mL) and xylazine (0.15µg/mL) and transcardially perfused with 2% paraformaldehyde. Brains were dissected and post-fixed for 30 minutes in the same fixative and cryoprotected in 30% sucrose/phosphate-buffered saline (PBS) before being flash-frozen in Optimal Cutting Temperature (OCT, Sakura) compound on dry ice and stored at –20°C.

Brain tissue from adult *Calcrl-*tdTomato mice was obtained from mice made by crossing *Calcrl*^*Cre*^ mice with a *Rosa26-flox-stop-tdTomato* reporter line, Ai14 (S. Han et al., 2015).

### RNAScope

Brains were sectioned (12 µm thick) and frozen sections were cryoprotected with OCT and mounted onto Superfrost Plus Slides (Thermo Fisher Scientific). We processed sections containing the insular cortex (AP Bregma: 0.02-0.26 mm) and quantified mRNA transcripts encoding the gene transcript for CGRP receptor (Calcrl and Ramp1) using RNAscope (Advanced Cell Diagnostics, Inc.) with integrated co-detection of Olig2 protein. According to manufacturer’s instructions, we performed tissue blocking and antigen retrieval prior to an overnight incubation at 4°C with rabbit anti-Olig2 antibody (Table 1). We then performed RNAscope hybridization using probes Mm-Calcrl-C2 (Cat No. 452281-C2) and Mm-Ramp1-O1-C3 (Cat No. 532681-C3) with fluorophore Opal 570 and 650 respectively (1:1000, Advanced Cell Diagnostics, Inc.) prior to a 30-minute secondary conjugation with Cy3-conjugated donkey anti-rabbit (Table 1) and counterstaining/mounting with ProLong™ Gold antifade mountant with DAPI (Invitrogen).

**Table 1.**
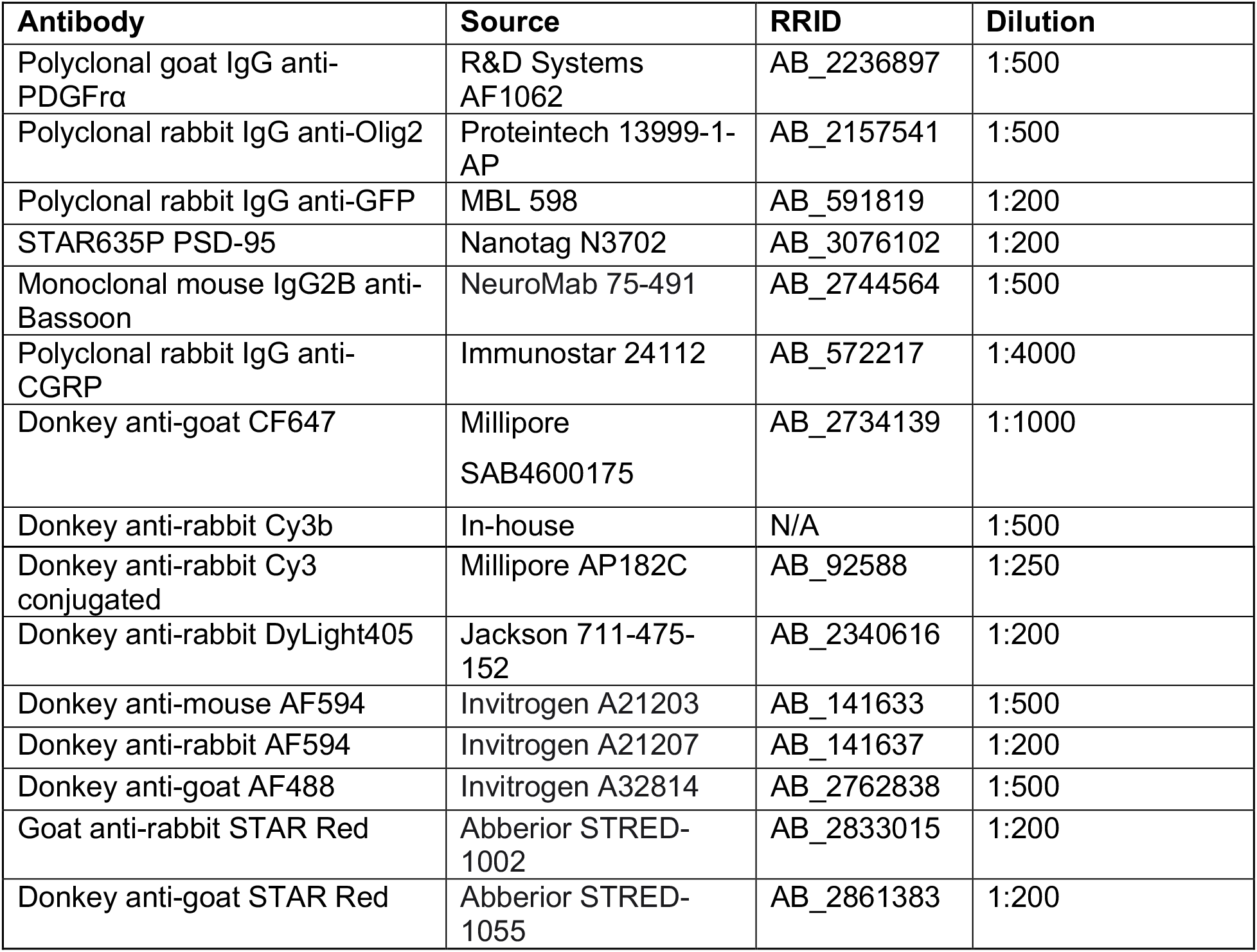
Primary and secondary antibodies used.

### Single-cell RNA-seq data processing

Single-cell RNA-seq data were obtained from a published study (Filipi et al., 2023) and processed using Seurat v5 R toolkit for single-cell genomics (Hao et al., 2024). Standard quality control and normalization were applied, including filtering cells with 200–6,000 detected genes and less than 10% mitochondrial content, log-normalization with a scale factor of 10,000, identification of 2,000 variable features, scaling of all genes, and dimensionality reduction using PCA and UMAP. Analysis was restricted to oligodendrocyte lineage cells under the control condition, using cell-type and condition annotations assigned in the original publication. This resulted in 82 OPCs, 42 committed OPCs (cOPC), and 2,371 oligodendrocytes (Oligo). Average expression of selected receptor genes was calculated per cell type, and genes with minimal expression in OPCs were excluded (average expression < 0.1).

### Virus Injection

We anesthetized the animals with isoflurane and placed them in a stereotaxic frame. Either left or right PB (−5.2 mm AP, ±1.5 mm ML, −2.9 mm DV) was targeted via a small craniotomy (∼1–2 mm). We injected 0.5 μl of adeno-associated virus generated by the University of Maryland School of Medicine’s Viral Vector Core, AAV5-DIO-ChR2-eYFP. Viruses were injected using a MICRO2T SMARTouch controller and Nanoliter202 injector head (World Precision Instruments) at a flow rate of 100 nl/min. The pipette was left in place for 10 min before being slowly retracted over 5–10 min. Mice were given Rimadyl for postoperative analgesia. Injection sites were verified by visually confirming robust eYFP fluorescence in the external PB.

### Immunohistochemistry

Brain sections were cut on a Leica CM1850 cryostat. For confocal experiments, 30 µm thick free-floating slices were kept in PBS for immunohistochemistry processing. For Stimulated Emission Depletion (STED) microscopy, 15 µm thick slices were directly mounted on 1.5H coverslips coated with (3-Aminopropyl)triethoxysilane (APTES) and air-dried.

Sections were blocked with 10% donkey serum (DS), 1% bovine serum albumin (BSA) and 0.3% Triton X-100 in PBS for 1h at room temperature. Sections were then incubated overnight at 4°C with primary antibodies diluted in 10% DS, 1% BSA, 0.1% Triton X-100 (Table 1). The following day, sections were rinsed 3x with PBS and incubated with secondary antibodies overnight (Table 1). The next day, sections were rinsed 3x with PBS. For confocal experiments, nuclei were stained with DAPI and sections were mounted with Prolong™ Gold antifade mountant. For sections imaged on the Abberior STED microscope, sections were mounted with Abberior Liquid Mount.

### Fluorescence Microscopy

Following RNAscope processing, we acquired images using a Leica microsystems TCS SP8 confocal microscope with a 40x oil-immersion objective. Z-stacks were created by sampling every 3 µm, with each image composed of 7-9 planes. Sections were reconstructed using Leica Las X Navigator tiling software.

CGRP receptor labeling images were also acquired using a Leica microsystems TCS SP8 confocal microscope. Tile scans were acquired using a 40x objective oil-immersion objective and z-stacks with a 1 µm step size in the insula, central amygdala (CeA), bed nucleus of the stria terminalis (BNST) and parabrachial nucleus (PBN) were acquired. For experiments in which distances between labeled OPC and CGRP processes were measured, 10 µm z-stacks were collected on a Nikon TI2 inverted microscope equipped with an Andor Dragonfly spinning disk confocal using a 60x oil-immersion objective and a z-step size of 0.3 µm. CGRP-OPC synapse images and Stimulated Emission Depletion (STED) images were acquired on the Abberior Facility Line STED microscope equipped with a pulsed 775nm STED laser using a 60x oil-immersion objective. From the tile scans, z-stacks were acquired with a step size of 200 nm for CGRP-OPC synapse images. 2D STED images were acquired with a pixel size of 30 nm x 30 nm for parabrachial CGRP (PBNcgrp) axon-OPC contacts, and 3D STED images were acquired with a pixel size of 50 nm x 50 nm x 250 nm to visualize CGRP within PBNcgrp axons.

In each experiment, excitation and detection parameters were held constant across all sections.

### Image Analysis

OPC *Calcrl* expression analysis was performed in ImageJ (version 1.54). 10 µm z-stacks were max-projected. Images were thresholded and converted into binaries, and the watershed tool was used to separate merged cells. Cells were segmented based on DAPI and cell-type specific marker expression. Cells were determined to express the CGRP receptor if their intensity values were >127.5 in the CGRP receptor (tdTomato) channel.

We used Imaris (version 10.2) to calculate CGRP-OPC distances. Images were preprocessed using a Gaussian filter and background subtraction tools. We created 3D reconstructions of OPCs and CGRP puncta using the surfaces tool for each channel. For each CGRP puncta, we identified the shortest distance to the OPC surface reconstruction. GraphPad Prism 10 was used to pool data and create distributions. Outliers were removed using Grubb’s test.

To identify regions of colocalization between OPCs, CGRP, and pre- and postsynaptic puncta, we assessed images processed using the Abberior TRUESHARP deconvolution tool using its default settings.

## Results

### Oligodendrocyte precursor cells express the receptor for calcitonin gene-related peptide

We sought to determine whether OPCs express the receptor for CGRP, using RNAscope and immunofluoresence. We performed RNAscope in the insular cortex, a cortical area of dense CGRP expression (Huang et al., 2021) using RNA probes for the CGRP receptor components receptor activity-modifying protein 1 (Ramp1) and calcitonin receptor-like receptor (Calcrl) (McLatchie et al., 1998). To identify oligodendroglia, we used an antibody for Olig2 (Table 1), a marker expressed throughout the oligodendrocyte lineage from OPCs to mature myelinating oligodendrocytes. In the insula, we identified cells that expressed Olig2 along with *Ramp1* and *Calcrl* (Fig. 1A and B). This suggested the presence of CGRP receptors in OPCs and/or oligodendrocytes. To determine if OPCs express CGRP receptors, we performed immunohistochemistry for platelet-derived growth factor receptor alpha (PDGFrα), an OPC-specific marker which extends throughout OPC processes and labels OPCs throughout the brain (Nishiyama et al., 2021). We focused on the insula as well as the parabrachial nucleus (PBN), central nucleus of the amygdala (CeA), and bed nucleus of the stria terminalis (BNST), regions of dense CGRP expression (Fig. 1C)(Huang et al., 2021). We used *Calcrl*-tdTomato mice, which display endogenous fluorescence (tdTomato) in cells which express *Calcrl*, an essential component of the CGRP receptor (S. Han et al., 2015). OPCs across the insula, BNST, CeA and PBN displayed CGRP receptor expression (Fig. 1D). We quantified the percentage of labeled OPCs that expressed *Calcrl* (Fig. 1E) and determined that 60% of OPCs in the insula and PBN, and 40% of OPCs in the BNST and CeA, expressed this receptor component (Fig. 1E). These findings provide the first demonstration that oligodendroglial lineage cells express components of the CGRP receptor, and that this expression is conserved across several forebrain and hindbrain regions.

**Figure 1.**
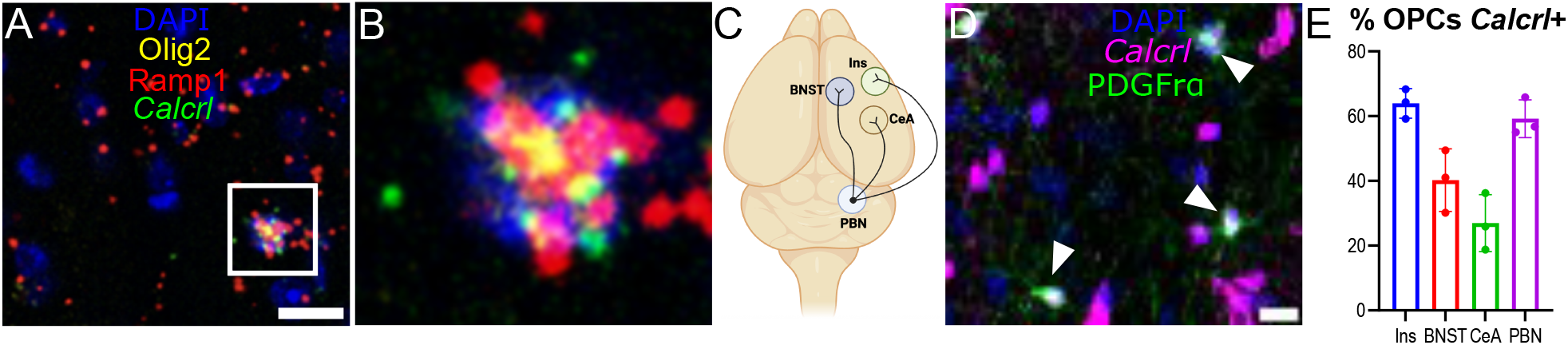
OPCs express the CGRP receptor. A) RNAscope images of Olig2+ cells in insula (yellow) expressing the CGRP receptor components *Calcrl* (green) and *Ramp1* (red). Scale bar = 20 μm. Higher magnification of region indicated by the white square is shown in B. C) Diagram showing several major efferent projection targets of parabrachial CGRP neurons. D) PDGFrα-expressing OPCs (green) expressing *Calcrl* (magenta; identified by expression of *Calcrl-*tdTomato; white arrowheads) in CeA. Similar overlap was observed in other brain regions. Scale bar = 20 µm. E) Percentage of *Calcrl+* OPCs in each brain region, defined by PDGFrα^+^ DAPI colocalized with *Calcrl* signal. Each data point represents data from one animal (2M, 1F).

Since RAMP1 antibodies are notoriously problematic for immunohistochemistry (Hendrikse et al., 2022), we used an orthogonal approach to further investigate this finding by analyzing an existing single-cell RNA-seq dataset to assess expression of CGRP receptor components in OPCs (Filipi et al., 2023), using Seurat v5 R toolkit for single-cell genomics (Hao et al., 2024).

From this dataset, we quantified the relative expression of *Calcrl* and *Ramp1* in OPCs versus committed OPCs (cOPCs) and mature oligodendrocytes (Oligos), as well as the relative expression of other neurotransmitter and G-protein coupled receptors (Fig. 2A). *Calcrl* and *Ramp1* displayed similar expression patterns to most other G-protein coupled receptors and neurotransmitter receptors across the oligodendroglia lineage, with expression widely peaking in the OPC stage and declining upon differentiation into oligodendrocytes (Fig. 2A). From this same dataset, we calculated the proportion of cells expressing both *Calcrl* and *Ramp1* across the oligodendroglial lineage (Fig. 2B). The majority of OPCs expressed both CGRP receptor subunits (60.97%, 50/82 cells), consistent with our histological estimates in the PBN and insula (Fig. 1E). In contrast, this proportion decreased across more differentiated populations: 16.67% of cOPCs (7/42 cells) and 0% (0/2371 cells) of oligodendrocytes expressed both receptor subunits. Consistent with this pattern, violin plots of *Calcrl* and *Ramp1* reveal robust expression in OPCs with a graded reduction across lineage progression (Fig. 2C). These data demonstrate robust OPC-specific expression of both CGRP receptor components in the mouse cortex.

**Figure 2.**
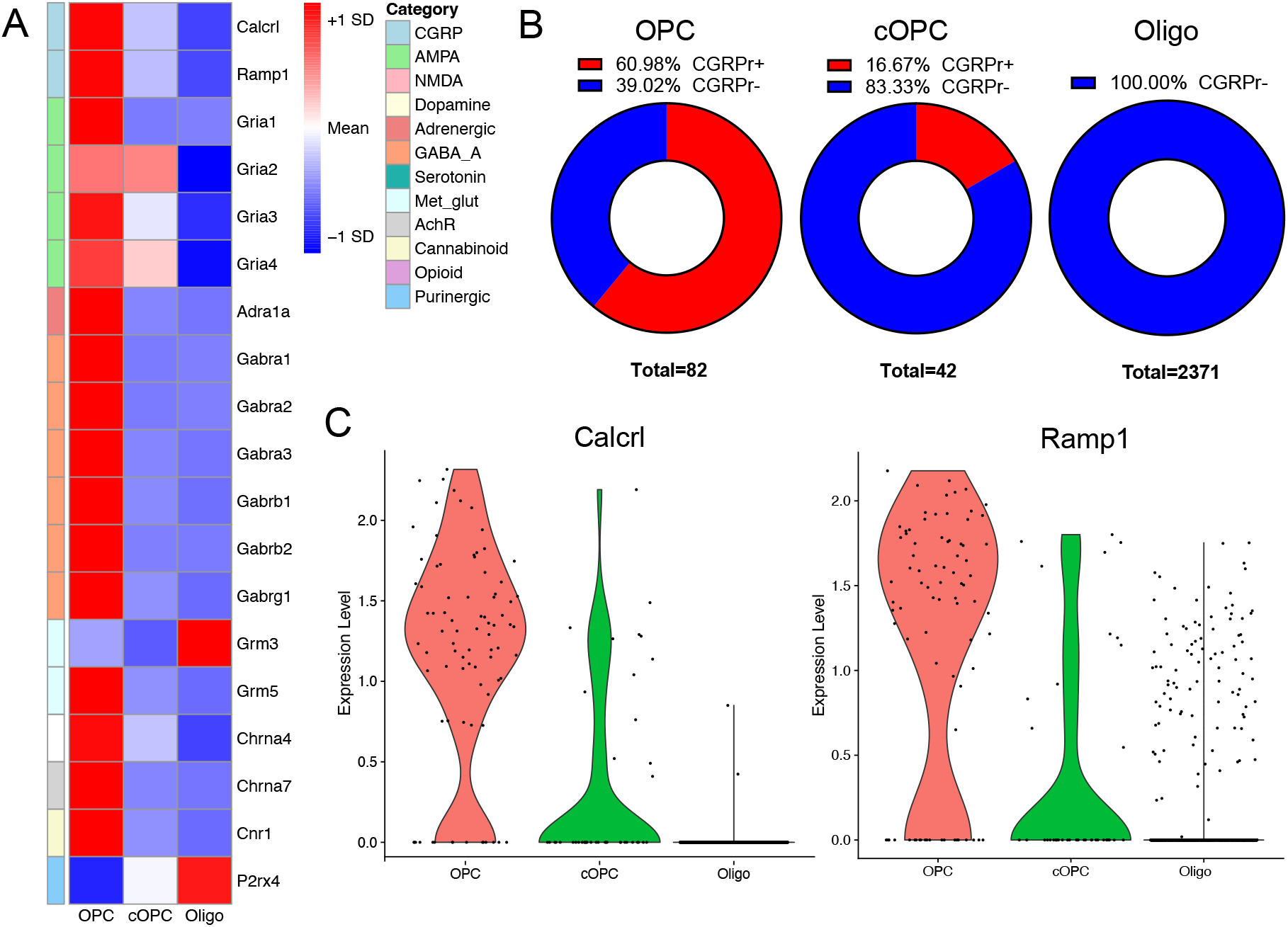
Single-cell datasets analysis of OPC CGRP receptor expression in mouse cortex. Analysis was performed on a previously published single-cell RNA sequencing dataset reported in Filipi et. al 2023. A) Heatmap showing the relative expression of *Calcrl* and *Ramp1* in OPCs versus committed OPCs (cOPC) and mature oligodendrocytes (Oligo), along with other G-protein coupled receptors and neurotransmitter receptors. Heatmaps display per-gene z-scored expression (row-wise), where red indicates higher expression relative to that gene’s mean across the lineage, blue indicates lower expression, and white indicates near-mean expression. B) Percentage of OPCs, cOPCs and Oligos which express both *Calcrl* and *Ramp1* C) Violin plots showing *Calcrl* and *Ramp1* expression in OPCs, cOPCs and Oligos.

### OPCs form close spatial relationships with CGRP axons

To determine if CGRP-containing axons contact OPCs, we examined the spatial proximity between OPCs and virally labeled CGRP-expressing efferents from the PBN (PBNcgrp axons). To identify PBNcgrp axons we injected AAV-DIO-eYFP unilaterally in the lateral PBN of *Calca-* cre mice. OPCs were identified with immunohistochemistry for PDGFrα. In each of the brain regions examined (PBN, CeA, BNST and insula) OPCs were abundant, their processes often intersecting with PBNcgrp axons (Fig. 3A). Super-resolution stimulated emission depletion (STED) microscopy confirmed the existence of close appositions between PBNcgrp axons and OPCs (Fig. 3B), with partial overlap of fluorescent labeling.

**Figure 3.**
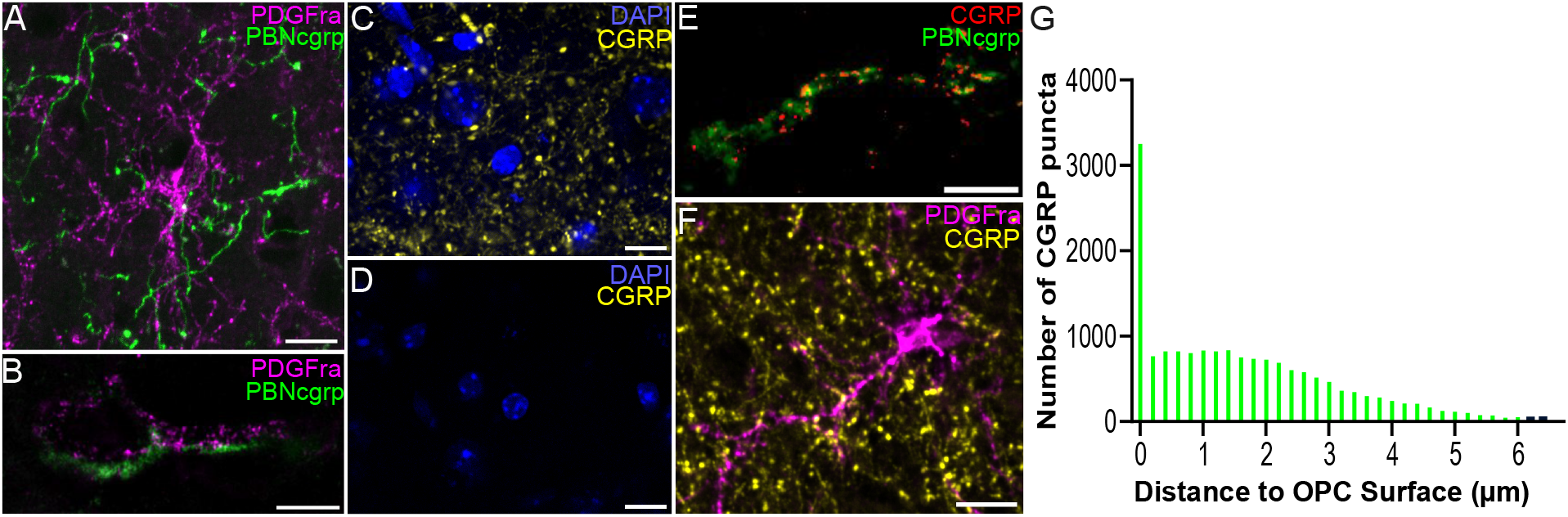
CGRP axons in close contact with OPCs. A) PBNcgrp axons virally labeled with cre-dependent eYFP (green) in CeA abutting PDGFrα-expressing OPC processes (magenta). Scale bar = 10 µm. B) High-resolution (STED) image of presumptive PBNcgrp neuron-OPC contacts in CeA. Scale bar = 2 µm. C) CGRP expression in CeA. Scale bar = 10 µm. D) Absence of CGRP labeling in CeA of homozygous *Calca-*cre transgenic mouse. Scale bar = 10 µm. E) STED image of CGRP (red) within virally labeled PBNcgrp axon (green) in BNST. Scale bar =2 µm. F) CGRP-containing axons (yellow) and OPCs (magenta) in CeA. Scale bar = 10 µm. G) Quantification of distance between CGRP puncta and OPCs. Data pooled from images of CeA, BNST and PBN, from three adult *Calca*-Cre mice (1M, 2F).

To precisely identify CGRP localization within PBNcgrp axons (as essentially all CGRP-expressing neurons in BNST and CeA come from PBN) (Shimada et al., 1985), we immunohistochemically labeled CGRP fibers. The CGRP antibody produced robust labeling in known regions of CGRP expression, including the CeA (Fig. 3C). No labeling was observed in the CeA of homozygous *Calca-*cre animals, which do not express CGRP, confirming the specificity of the CGRP antibody (Fig. 3D). STED microscopy revealed CGRP labeling in virally labeled PBNcgrp efferents, with CGRP puncta around 80-100nm in diameter, congruent with the size of large dense core peptidergic vesicles (Merighi, 2018) (Fig. 3E). CGRP puncta were often seen near PDGFrα+ OPC processes (Fig. 3F). We created 3D reconstructions of CGRP puncta and OPC surfaces in Imaris™ to measure the nearest distance of each puncta to an OPC. Over 40% of CGRP puncta were observed within 1µm or less of an OPC, and 18% of CGRP puncta colocalized with OPC labeling (Fig. 3G). Together, these data demonstrate the close spatial proximity between CGRP and OPCs in several brain regions, with a high degree of colocalization suggesting the potential presence of synaptic contacts.

### CGRP is observed at putative synaptic contacts with OPCs

To determine whether close contacts between CGRP and OPCs may represent synaptic sites, we determined if pre- and postsynaptic markers are located at sites of interactions between CGRP and OPCs (Fig. 4A). PBNcgrp neurons co-release glutamate (Lorsung et al., 2025; Palmiter, 2018); therefore, we selected the postsynaptic protein postsynaptic density-95 (PSD-95), a scaffolding protein involved in localizing glutamate receptors to the plasma membrane. Transcriptomic datasets report that PSD-95 is expressed by OPCs (Marques et al., 2016; Y. Zhang et al., 2014). Further, in zebrafish, PSD-95 is expressed throughout the brain in excitatory synapses with OPCs (Li et. al 2024). Consistent with this finding, we identified PSD-95 in OPC processes in mice (Fig. 4B).

**Figure 4.**
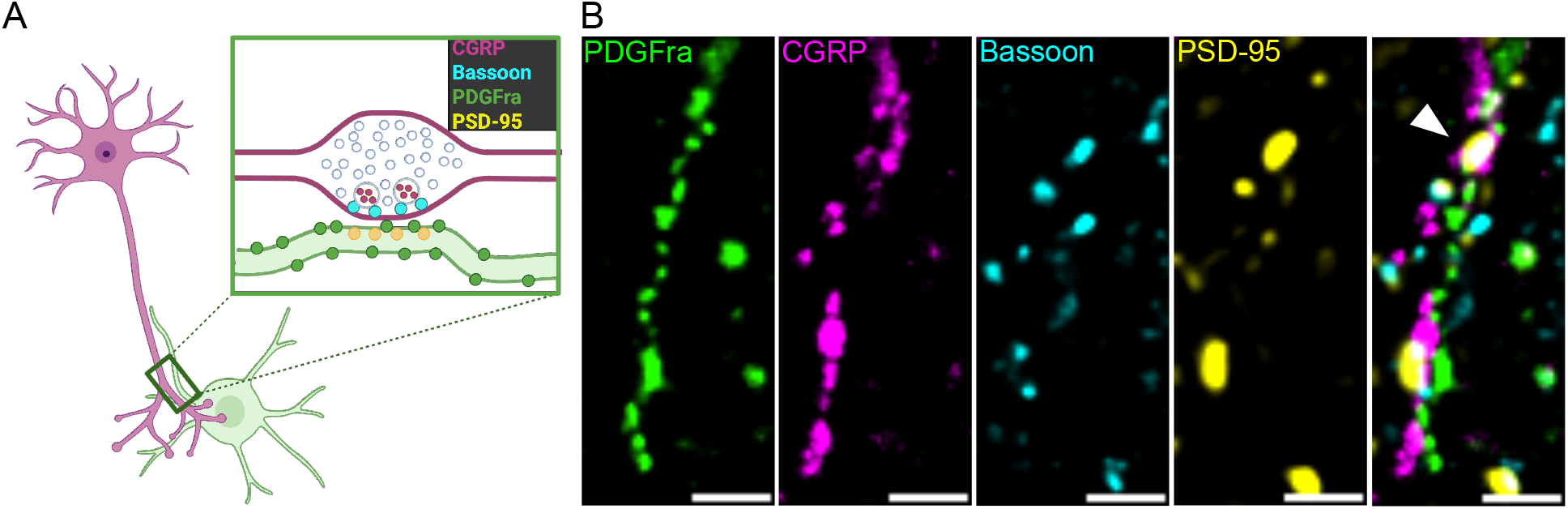
CGRP can localize to putative neuron-OPC synapses. A) Hypothesis schematic showing predicted phenomenon and targets labeled with immunohistochemistry. Created with Biorender.com B) Colocalization of PDGFrα, CGRP, Bassoon and PSD-95 in mouse PBN (indicated by white arrowhead). Scale bar = 1 µm.

To identify presynaptic sites, we selected an antibody to Bassoon, a protein involved in presynaptic active zone assembly. Bassoon is present at virtually all excitatory and inhibitory synapses in the CNS (Südhof, 2012; tom Dieck et al., 1998). We performed these experiments using immunohistochemistry for PDGFrα to label OPC somas and processes. We used CGRP and OPC channels from tile scans of the PBN to identify regions of CGRP–OPC apposition for z-stack acquisition. At these sites of interest, we collected z-stacks of CGRP, PDGFrα, and pre-and postsynaptic markers Bassoon and PSD-95 to assess the presence of putative neuron-OPC synapses, defined by colocalization of OPCs with synaptic puncta. Bassoon and PSD-95 colocalized extensively in areas of the PBN distant from PDGFRα+ processes, as expected given the abundance of neuronal synapses. CGRP immunoreactivity was occasionally colocalized at these presumptive neuronal synapses. However, we also found examples of CGRP colocalizing with Bassoon and PSD-95 at sites of OPC contact (Fig. 4B). Upon qualitative examination, these colocalized synaptic puncta at OPCs did not appear different in size nor intensity from those not associated with PDGFrα. These data are consistent with the presence of CGRP at synapses onto OPCs.

## Discussion

We characterized relationships between axons expressing calcitonin gene-related peptide (CGRP) and oligodendrocyte precursor cells (OPCs) in the mouse brain. We focused on the parabrachial nucleus (PBN), a main source of CGRP in the brain (Palmiter, 2018), and its efferent projection targets. We show that OPCs express the CGRP receptor and form close contacts with CGRP-containing axons, some of which may represent neuron-OPC synapses. To our knowledge, this is the first demonstration of the necessary cellular components of CGRP signaling in OPCs.

### OPCs express the CGRP receptor

Our understanding of CGRP’s effects in the brain has previously been restricted to its actions on neurons, which express CGRP receptor in cortical and subcortical regions (Warfvinge & Edvinsson, 2019). We provide the first demonstration that mammalian OPCs in the PBN and several of its major targets, including the central amygdala, bed nucleus of the stria terminalis (BNST), and insular cortex, express both *Calcrl* and *Ramp1*, necessary components of the CGRP receptor (McLatchie et al., 1998). This finding is consistent with single-cell sequencing studies that demonstrate OPC expression of CGRP receptor components in the mouse cortex (Filipi et al., 2023). These data position OPCs as a novel cell type which may directly respond to CGRP input.

### Potential modes of CGRP signaling to OPCs

Nearly half of labeled CGRP puncta were within 1μm of a process labeled for platelet-derived growth factor receptor alpha (PDGFRα), a marker for OPCs (Nishiyama et al., 1999). Labeled CGRP puncta were present along axons (presumably originating from PBN), and the majority were <5 μm from an OPC, well within the expected diffusion distance of peptides in extracellular space (van den Pol, 2012). These data provide evidence for anatomical substrates for CGRP signaling to OPCs.

Like other neuropeptides, CGRP is packaged in large dense-core vesicles (LDCVs) (Merighi et al., 1991). Neuropeptide signaling may occur as volumetric, bulk transmission, often at non-synaptic sites (Fuxe et al., 2007; Jansson et al., 2002; van den Pol, 2012). However, in the hippocampus, LDCVs can colocalize at conventional chemical synapses, sites of fast neurotransmitter release mediated by small synaptic vesicles, suggesting that some neuropeptide signaling can occur at these synapses (Lochner et al., 2006; van de Bospoort et al., 2012). Indeed, our data support the localization of CGRP-containing puncta (presumably LDCVs) at conventional synaptic release sites onto OPCs (Fig. 4B). This suggests that CGRP can be released at chemical synapses with OPCs.

However, the anatomical approaches we used cannot conclusively identify CGRP release sites. We rely on an indirect approach, using the presence of synaptic markers of chemical synapses. We label the presynaptic active zone with Bassoon, and the postsynaptic zone with PSD-95, to identify presumptive synapses onto OPCs. OPCs express PSD-95 (Marques et al., 2016; Y. Zhang et al., 2014) and it has been used as a postsynaptic marker of neuron-OPC synapses in zebrafish (J. Li et al., 2024), where it may serve to anchor AMPA receptors at the postsynaptic membrane (Chen et al., 2015; Ehrlich & Malinow, 2004). We emphasize that there is currently no direct evidence that either Bassoon or PSD-95 are involved in LDCV release.

There are currently no markers for LDCV release sites, and there is little evidence of specific proteins that may be uniquely required for LDCV release. For example, the vesicle priming protein, Munc-13, required for synaptic vesicle release (Augustin et al., 1999; Varoqueaux et al., 2002) appears to be involved in controlling the location and efficiency of LDCV release, but is not essential for LDCV release (van de Bospoort et al., 2012). Therefore, the immunohistological tools do not currently exist to determine whether CGRP is primed for release, or simply located nearby, either extrasynaptic or synaptic release sites (i.e classical chemical synapses). Our findings suggest that, in the PBN, CGRP is at least present at release sites for classical chemical synapses, though we cannot rule out the possibility that CGRP may also be released extrasynaptically.

Our data suggest both direct synaptic CGRP signaling to OPCs, supported by CGRP at presumptive chemical synapses onto OPCs, and extrasynaptic signaling, indicated by close apposition of CGRP with OPCs outside of these sites. Distinguishing between these potential modes of transmission will require localization of the CGRP receptor within OPCs.

Unfortunately, there are currently no reliable commercial antibodies for either CGRP receptor component – Calcrl or Ramp1 (Hendrikse et al., 2022). Ongoing developments in multiplexed nanoimaging should, in the future, help determine the spatial and cellular organization comprising the molecular basis for CGRP-OPC communication.

### Potential outcomes of CGRP signaling to OPCs

Neuropeptides may modulate the activity of neurotransmitters they are released with, to influence the efficacy of synaptic signaling (Russo, 2017; Tallent, 2008). PBN afferents co-release glutamate and CGRP, and we and others have shown that CGRP can potentiate the effects of glutamate in neurons (Campos et al., 2018; J. S. Han et al., 2010; Lorsung et al., 2025; Okutsu et al., 2017). Whether CGRP may similarly influence glutamatergic signaling to OPCs is an important open question, because synaptic activity to OPCs is thought to control a range of their functions including differentiation and myelination. For instance, in zebrafish, modulation of glutamate synapses onto OPCs is shown to influence OPC lineage progression and myelination (J. Li et al., 2024). Therefore, CGRP-mediated potentiation of glutamatergic signaling to OPCs may constitute a mechanism through which CGRP influences OPC behaviors.

CGRP might influence OPC behaviors, including proliferation and differentiation, through mechanisms independent from glutamatergic regulation. It is useful to consider another peptide involved in pain pathogenesis, pituitary adenylate-cyclase activating peptide (PACAP), (Davis-Taber et al., 2008; Markovics et al., 2012; Narita et al., 1996), which promotes OPC proliferation through cAMP (Lee et al., 2001; Lelievre et al., 2006). The cAMP/PKA pathway has also been implicated downstream of the CGRP receptor in neurons (J. S. Han et al., 2010; Kunioku et al., 2023), and both cAMP/PKA and PKC signaling influence OPC proliferation and differentiation (Althaus et al., 1991; Baron et al., 1999; Hoi et al., 2023; Joubert et al., 2010). Thus CGRP signaling could hypothetically control OPC proliferation or differentiation through these pathways.

CGRP input to OPCs may also influence other OPC functions, such as immune modulation, regulation of the blood-brain barrier (BBB), or synapse engulfment (Buchanan et al., 2023; Fang & Bai, 2023). CGRP plays a neuroprotective role through BBB modulation (Liu et al., 2011; Xiong et al., 2023), and regulates immune response through functions such as regulation of lymphocyte differentiation and cytokine production (Assas et al., 2014). It will be important to establish whether OPC activation contributes to these established functions of CGRP.

### CGRP, OPCs and chronic pain

Both OPCs and CGRP are associated with chronic pain and stress conditions. CGRP is causally related to several conditions, particularly migraine and cluster headache disorders (Dussor, 2019; Ho et al., 2010; Iyengar et al., 2017). There is growing evidence for the role of OPCs in particular, and oligodendroglia in general, in chronic pain (Kim & Angulo, 2025). Pain following sciatic nerve injury is associated with an increase in proliferating OPCs in the spinal cord (Echeverry et al., 2008), and the number of cortical OPCs increases after spinal cord injury (C. Li et al., 2022). These changes may relate to downstream abnormalities in myelination associated with a number of chronic pain conditions (Arkink et al., 2017; DaSilva et al., 2007; Mansour et al., 2013; Palm-Meinders et al., 2017). Understanding whether and how CGRP signaling in OPCs is involved in the pathogenesis of these conditions may aid therapeutic development.

## Acknowledgements

We thank Dr. Yarimar Carrasquillo (National Center for Complementary and Integrative Health, National Institutes of Health, Bethesda, Maryland) for providing the tissue from the *Calcrl*-tdTomato mice.

## Notes

### Competing Interest Statement

The authors have declared no competing interest.

